# MyD88 Mediates Colitis- and RANKL-induced Microfold Cell Differentiation

**DOI:** 10.1101/2021.09.16.460646

**Authors:** Yang Li, Shanshan Yang, Xin Huang, Ning Yang, Caiying Wang, Jing Zhao, Zhizhong Jing, Luc Willems, Guangliang Liu

**Author notes:** These authors contributed equally to this work (Yang Li, Shanshan Yang). Corresponding author: Guangliang Liu, 1 XuJiaPing, YanChangBu, ChengGuan District, Lanzhou, Gansu 730046, China.

## Abstract

Intestinal microfold (M) cells are critical for sampling antigen in the gut and initiating the intestinal mucosal immune response. In this study, we found that the differentiation efficiency of M cells was closely related to the colitis severity. The expression levels of M cells differentiation-related genes were synchronized with the kinetics of pro-inflammatory cytokines expression originated from dextran sulfate sodium (DSS) induction and *Salmonella* infection. Compared with wild-type (WT) mice, *MyD88*^-/-^ mice exhibited significantly lower expression levels of M cells differentiation-related genes. However, DSS could induce colitis in *MyD88*^-/-^ mice but failed to promote M cells differentiation. Furthermore, the receptor activator of the Nuclear Factor-κB ligand (RANKL) induced M cells differentiation in murine intestinal organoids prepared from both WT and *MyD88*^-/-^ mice. However, less M cells differentiation were found in *MyD88*^-/-^ mice as compared with WT mice. Hence, we concluded that myeloid differentiation factor 88 (MyD88) is an essential molecule for colitis- and RANKL-related M cells differentiation.

## Introduction

M cells are specialized epithelial cells located mainly at follicle-associated epithelium (FAE) [1] and villous epithelium [2] of the small intestines. These cells are also found at the epithelium of colonic mucosa [3] and nasopharynx-associated lymphoid tissue (NALT) [4]. Although M cells are not antigen-presenting cells (APC), they play essential roles in mucosal immune surveillance [5]. M cells possess unique morphological features, such as irregular brush border, pocket structure, reduced glycocalyx, and microvilli [6]. They are conventionally known for capturing luminal antigens and delivering antigens particles to underlying immune cells. Many intestinal pathogens, such as murine Norovirus [7, 8], Reovirus [8], *Salmonella typhimurium* (*S. Typhimurium*) [9], and *Candida albicans* (*C. albicans*)[10] utilize M cells as the entry for invasions.

M cells are differentiated from leucine-rich repeat-containing G-protein coupled receptor 5^+^ (Lgr5^+^) stem cells [11]. In this differentiation process, the receptor activator of the Nuclear Factor-κB ligand (RANKL) is indispensable. Enterocytes expressing receptor activator of NF-κB (RANK) are stimulated by RANKL, then differentiated into M cells [12]. Spi-B, a transcription factor of the Est family, also mediates this process [11, 13]. Some gut microbiota impact M cells differentiation as well. For example, bacterial flagellin recovers the age-related decline of FAE M cells in mice [14]. The number of M cells in specific pathogen-free (SPF) mice increases after 7-day feeding in a typical animal house environment. *Salmonella enterica* serovar Typhimurium type III could also increase the number of M cells[15]. Additionally, tumor necrosis factor-α (TNF-α), an inflammatory cytokine, induces M cells differentiation in the colon [3]. TNF-α is induced by MyD88 or tumor necrosis factor receptor-associated factor 6 (TRAF6) through stimulating the production of NF-κB [16, 17]. As the downstream molecule of RANKL-RANK signaling, TRAF6-mediated Nuclear Factor-κB (NF-κB) signaling pathway is responsible for developing M cells [18]. Therefore, we hypothesized that MyD88 might participate in colonic M cells differentiation.

In this study, we show that DSS and *Salmonella choleraesuis* (*S. choleraesuis*) cause severe colitis in mice. M cells differentiation is attenuated when MyD88 is knocked out. Meanwhile, DSS-induced colitis promotes glycoprotein 2^+^ (GP2^+^) M cells production in WT mice but not in *MyD88*^-/-^ mice. Furthermore, RANKL upregulates M cell-specific gene expression on intestinal organoids partly dependent on MyD88. Therefore, we concluded that MyD88 mediates colitis- and RANKL-induced colitis-related colonic M cells differentiation.

## Methods and Materials

### Animals and related materials

*MyD88*^-/-^ (B6.129P2(SJL)-*MyD88*^tm1.1Defr^/J, Stock No: 009088) mice were originally purchased from The Jackson Laboratory (Anthony L. DeFranco, University of California, San Francisco). All mice used in this study were 6-8 weeks old and were bred in the SPF facility of Lanzhou Veterinary Research Institute (LVRI). Dextran sodium sulfate (DSS) was purchased from HWRK Chemical Company. *S. choleraesuis* (Strain NO. CVCC79102) was purchased from the China Institute of Veterinary Drug Control. SPF WT (wild type) C57BL/6 mice were purchased from the Experimental Animal Center of LVRI. All experimental procedures and animal care protocols were carried out by following Care and Use of Laboratory Animals of LVRI, Chinese Academy of Agricultural Sciences, China.

### Inflammatory models

For the DSS-induced inflammatory model, DSS 3% (M.W 40000) was added to the drinking water for the free intake for seven days. DSS-free drinking water was used as a control for the mice in the control group. For the *S. choleraesuis*-induced inflammatory model, *S. choleraesuis* were collected by centrifugation and re-suspended in PBS buffer. The mice in the infected group were gavaged with 10^4^ CFU of *S. choleraesuis* or with PBS as control. All mice were kept in the same environment, weighed daily and sacrificed at day 8.

### Histological analysis

After euthanasia, mice colons were fixed in the FinePix tissue-fix solution (RightTech, China) for 24h, dehydrated according to the standard protocol, and then embedded in paraffin. Then tissue blocks were sectioned and deparaffinized in xylene, stained with Haematoxylin and Eosin (H&E). Finally, double-blinded histological analysis was performed to evaluate the colitis level as described [19].

### Western Blot analysis

Total proteins were prepared from tissue samples of the colon using RIPA lysis buffer (Beyotime, China) with protease inhibitor. A total of 20 μg of protein were loaded in each well for SDS-PAGE, transferred, and immunoblotted overnight at 4°C with the primary antibody against GP2 (1:1000, MBL D278-3, Japan) and GAPDH (1:5000, Proteintech Group, U.S.). Goat anti-Rat IgG (OriGene Technologies, U.S.) and anti-Rab IgG secondary rabbit antibody (OriGene Technologies, U.S.) were diluted at 1:5000 for incubation with the membrane. Finally, the immunoblots were developed with chemiluminescence detection reagents (Advansta, U.S.)[20, 21].

### Murine intestinal organoids culture

Mice were sacrificed to harvest the small intestines. After being washed by PBS buffer, the intestines were cut into pieces and dissociated with 2 mM EDTA buffer in 15 mL centrifuge tubes for 20 min at 4°C. The samples were centrifuged at 300 ×g for 5 min to remove the EDTA buffer, repeatedly pipetted with 5 mL PBS buffer. The crypts were then collected through a 70 μm strainer (BD, U.S.), centrifuged at 300 ×g for 5 min, and cultured in the 24-well tissue culture plate (Corning, U.S.) with Matrigel Matrix (Corning, U.S.) and Murine Intestinal Organoids Growth Medium (STEMCELL, Canada). For the RANKL stimulation assay, organoids were stimulated with 100 ng/mL recombinant human RANKL (CST, U.S.) for one day (PBS buffer was used as control). The RANKL and mock control reagents were added daily.

### RNA extraction and quantitative real-time PCR

After the mice were sacrificed, their colon was washed by cold PBS then harvested. The total RNA was extracted from the colon samples using RNAiso reagent (TaKaRa, Japan) and used for cDNA preparation using Honor II 1st Strand cDNA Synthesis SuperMix (Novogene, China) with hexamer random primers. To evaluate the inflammation responses and M cells differentiation within mouse colon, the relative expression levels of TNF-α, IL-1β, IL-6, GP2, Spi-B, RANK, TNF-α-induced protein 2 (Tnfaip2), and C-C motif ligand 9 (CCL9) were determined by RT-qPCR using ChamQ SYBR qPCR Master Mix (Vazyme, China). The primers used in this study are listed in Table 1.

**Table 1.**
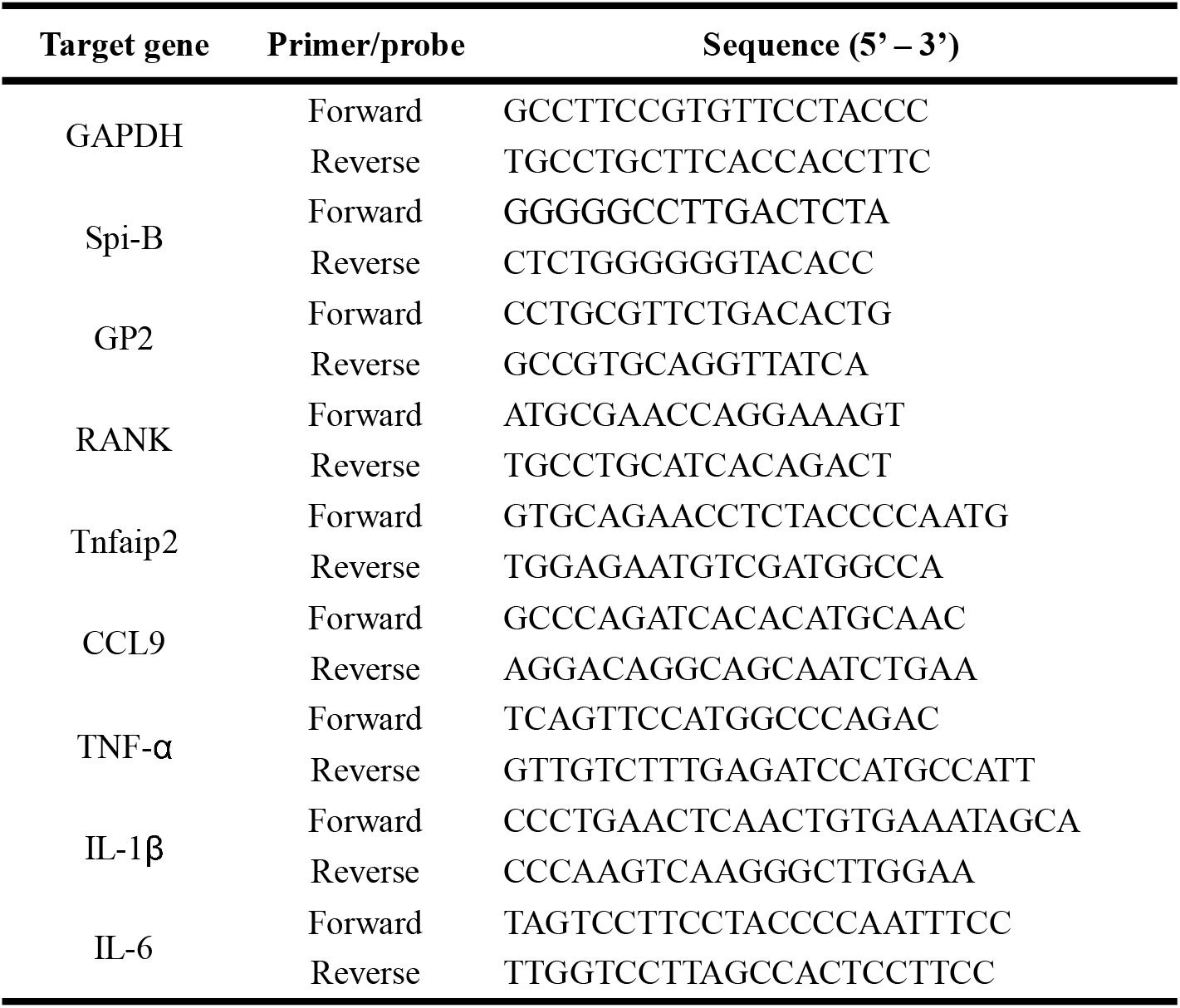
Primers used in this study

### Statistical analysis

The data were presented as the means ± SEM. The significance between groups was analyzed by one-way analysis of variance or Student’s *t*-test with GraphPad Prism 7 software and *P* values.

## Results

### DSS induces colitis and GP2^+^ M cells differentiation

To investigate the relationship between colonic M cells differentiation and colitis, a model of DSS-induced colitis was used. After oral administration of DSS for 7 days, the DSS-treated mice lost weight compared with the control group (Figure 1A). The spleen weight increased in DSS-treated mice, but their colon weight and length were reduced (Figure 1B). The mRNA levels of pro-inflammatory cytokine TNF-α, IL-1β, and IL-6 in the colon were quantified by RT-qPCR. DSS-treated mice had significant-high mRNA levels of TNF-α, IL-1β, and IL-6 (Figure 1C). Histological changes, including inflammatory cell infiltration in the basal layer and crypt, edema and reduced mucus, were observed in the colon of DSS-treated mice (Figure 1D). To assess the differentiation of colonic M cells, the mRNA level of M cells differentiation-related genes, including GP2, Spi-B, RANK, Tnfaip2, and CCL9, were measured by RT-qPCR. The results showed that the mRNA levels of GP2, Tnfaip2, and CCL9 from the colon of the DSS-treated group were significantly upregulated compared to - control - (Figure 1E). The immunoblotting results showed that DSS treatment promoted the expression of GP2 (Figure 1F). These results indicated that oral administration with DSS successfully induced colitis in mice. In addition, RT-qPCR suggested that DSS treatment induced GP2^+^ M-cell differentiation.

**Figure. 1.**
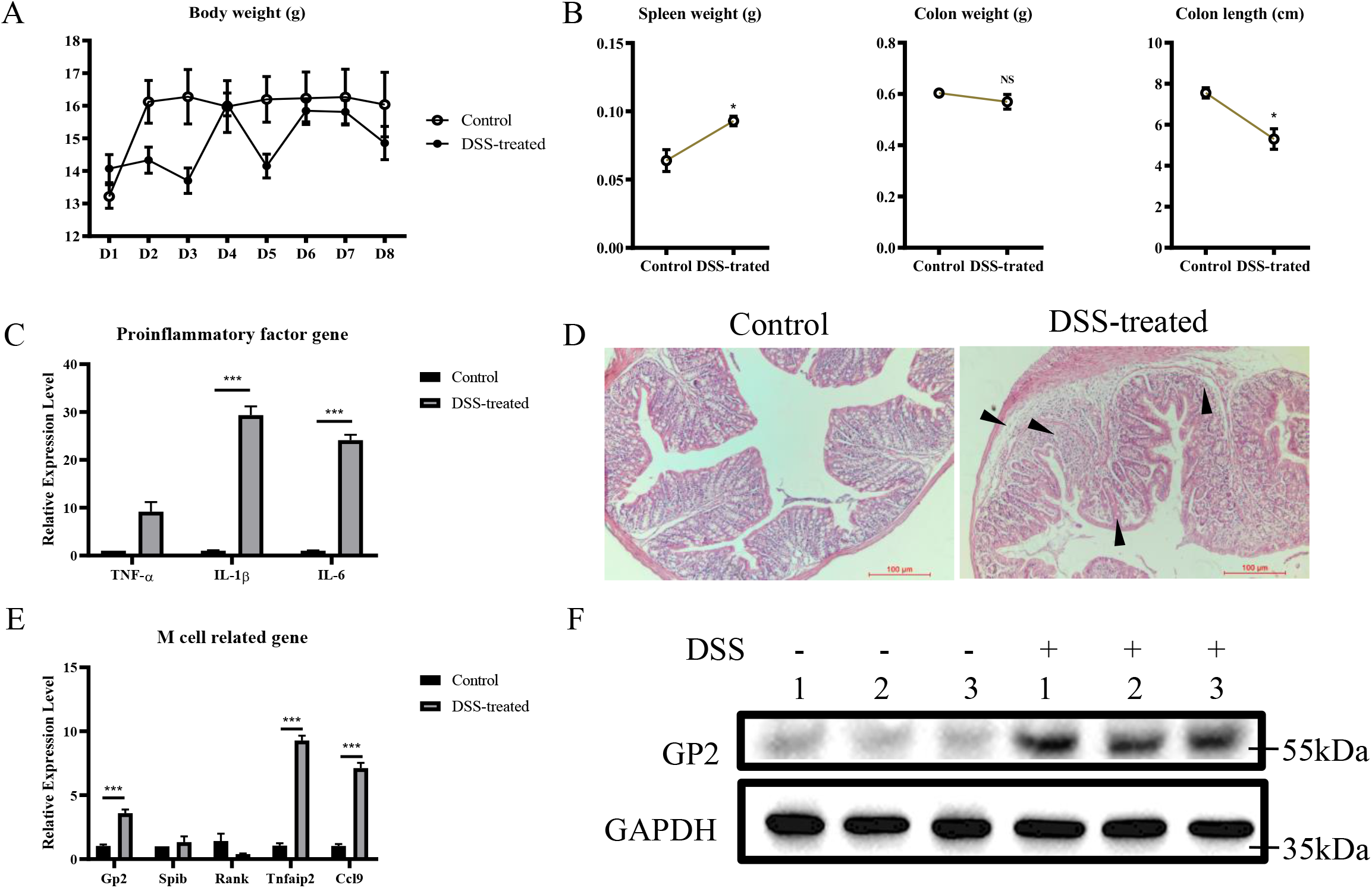
DSS induces colitis and GP2^+^ M cells differentiation. WT mice in the DSS-treated group (n=3) and control group (n=3) were allowed to intake DSS-contained or normal water for 7 days freely. Their body weights were monitored daily (A). The mice were sacrificed on day 8, and their spleen weight, colon weight, and colon length were measured for assessing pathological changes caused by DSS treatment (B). Total RNA from colon samples was used to detect the mRNA levels of TNF-α, IL-1β, IL-6 by RT-qPCR (C). Partial colon samples were used for histological analysis to evaluate the colitis (D). Total RNA from colon samples was used to detect the mRNA levels of GP2, Spi-B, RANK, Tnfaip2, and CCL9 by RT-qPCR (E). The data were calculated using the 2^-ΔΔCt^ method. *, *P*<0.05; **, *P*<0.01; ***, *P*<0.001. The GP2 expression levels in total protein from colon samples were evaluated by Western blot (F).

### *S. choleraesuis* infection induces colitis and GP2^+^ M cells differentiation

To further verify that colitis is related to colonic M cells differentiation, mice were administrated with *S. choleraesuis* orally. Mice in the *S. choleraesuis*-treated group suffered weight loss compared to the controls (Figure 2A). Typical symptoms, such as the increase in spleen weight, and a reduction of colon weight and length, were observed in the S. choleraesuis-treated group (Figure 2B). Concomitantly, bacterial colonized the liver and spleen (Figure 2C). The mRNA levels of TNF-α, IL-1β, and IL-6 in the colon were detected by RT-qPCR. The results illustrated that the *S. choleraesuis* infection significantly upregulated the mRNA levels of TNF-α and IL-6 can in mouse colons (Figure 2D). Pathological changes, including inflammatory cell infiltration in the basal layer and crypt, fibrinous exudation and reduced mucus, indicated that the *S. choleraesuis* induced colitis (Figure 2F).

**Figure. 2.**
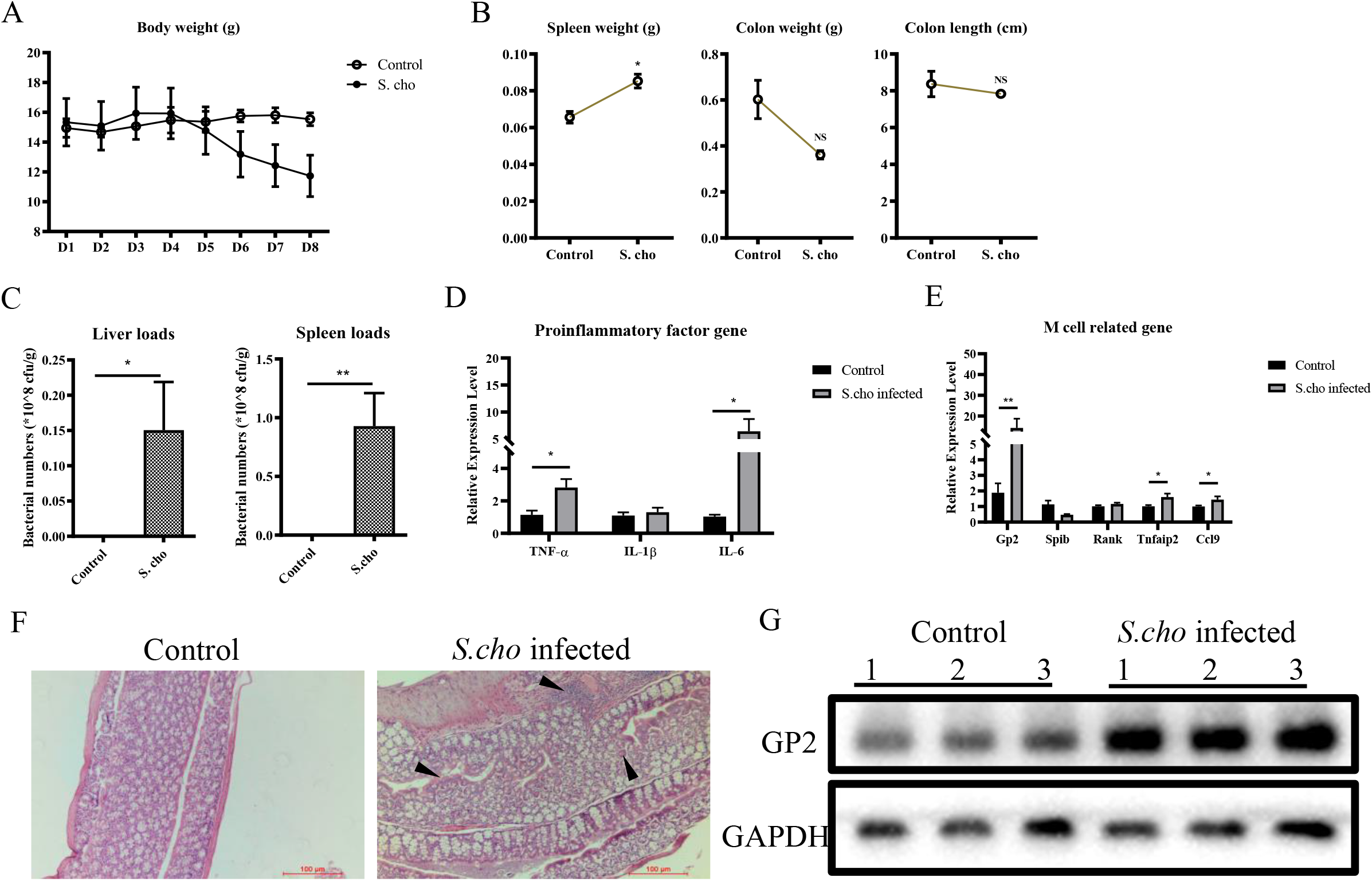
*S. choleraesuis* infection induces colitis and GP2^+^ M cells differentiation. WT mice were gavaged with *S. choleraesuis*-PBS buffer mix (*S. choleraesuis*-infected group, n=3) or normal PBS buffer (control group, n=3). Their body weight was monitored daily (A). The mice were sacrificed on day 8, and the spleen weight, colon weight, and colon length were evaluated to assess the pathological changes caused by *S. choleraesuis* infection (B). The bacterial loads in the liver, spleen were analyzed (C). Total RNA from colon samples was used to detect the mRNA level of TNF-α, IL-1β, IL-6 by RT-qPCR (D). Total RNA from colon samples was used to detect the mRNA levels of GP2, Spi-B, RANK, Tnfaip2, and CCL9 by RT-qPCR (E). Tissue samples from the colon were used for histological analysis for assessing colitis (F). The data were calculated using the 2^-ΔΔCt^ method. *, *P*<0.05; **, *P*<0.01; ***, *P*<0.001. The GP2 expression levels in total protein from colon samples were evaluated by Western-blot (G).

To evaluate the M cells differentiation, the mRNA levels of GP2, Spi-B, RANK, Tnfaip2, and CCL9 were measured by RT-qPCR. The result showed that the mRNA levels of GP2, Tnfaip2, and CCL9 in the colon from the *S. choleraesuis*-infected group were significantly higher than those in the mock-control group (Figure 2E). Western-blot analysis confirmed that *S. choleraesuis* infection induced the GP2 expression (Figure 2G). Taken together, these data demonstrate that *S. choleraesuis* infection induces colitis and GP2^+^ M cells differentiation

### MyD88 is a critical factor for colonic M cells differentiation

We speculated that MyD88 might play a role in the differentiation of murine colonic M cells. To test this hypothesis, we compared WT and *MyD88*^-/-^ mice. During 7-day housing in the same environment, the bodyweight of both *MyD88*^-/-^ and WT mice remained stable (Figure 3A). On day 7, the spleen weight of *MyD88*^-/-^ mice were lower than that of WT mice, but their colon weight and length were not significantly different (Figure 3B). Histological analysis of the colons did not reveal differences between *MyD88*^-/-^ and WT mice (Figure 3C). To investigate whether *MyD88*^-/-^ affects the differentiation of colonic M cells, mRNA level of M cells differentiation-related genes was detected by RT-qPCR while the GP2 protein expression was measured by western-blot. The results showed that the *MyD88*^-/-^ mice had significantly lower mRNA levels of GP2 and Spi-B, as well as GP2 protein levels compared to WT mice (Figure 3D, 3E). These results indicated that the lack of MyD88 restrained the differentiation of colonic M cells and suggested that the MyD88 is a critical factor during the colonic M cells differentiation.

**Figure. 3.**
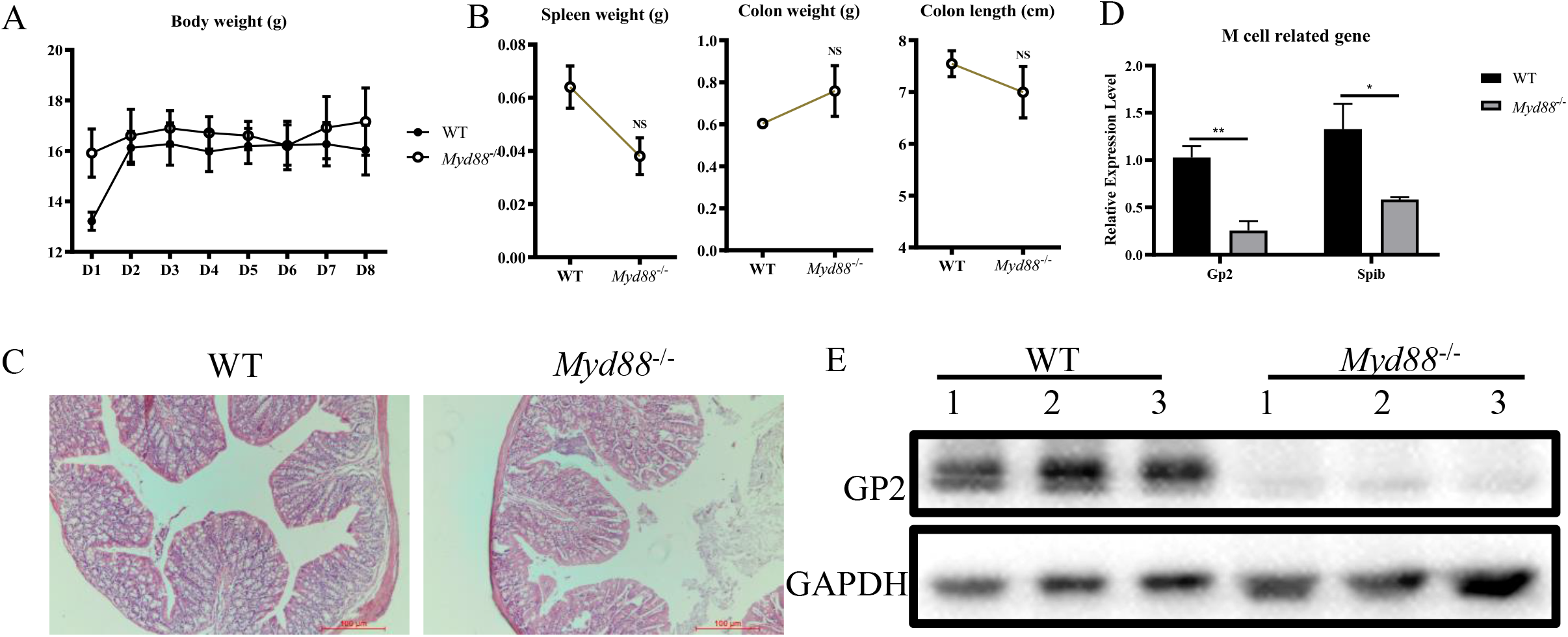
MyD88 is a critical factor for colonic M cells differentiation. The body weight of WT and *MyD88*^-/-^ mice (n=3) were monitored for 8 days (A). After sacrificing, their spleen weight, colon weight, and colon length were measured (B). Partial colon samples were fixed for histological analysis (C). Total RNA from colon samples was used to detect the mRNA levels of GP2 and Spi-B by RT-qPCR (D). The GP2 expression levels in complete protein from colon samples were evaluated by Western-blot (E).

### MyD88 is involved in colitis induced M cells differentiation

To explore whether MyD88 participates in colitis-induced M cells differentiation, *MyD88*^-/-^ mice were given DSS orally for 7days. Compared to the control group, weight loss was observed in the DSS-treated mice (Figure 4A). Moreover, the DSS treatment group was associated with more severe clinical symptoms, characterized by increased spleen weight, decreased colon weight, and colon length (Figure 4B). Pathological changes, including inflammatory cell infiltration, exfoliation of epithelial cells, crypt branching and decreased mucus (Figure 4C) and the higher mRNA levels of TNF-α, IL-1β, and IL-6 in the colon of the DSS-treated group (Figure 4D), indicated that DSS still induced colitis in *MyD88*^-/-^ mice. However, during this severe inflammatory response, M cells differentiation-related markers decreased (GP2, Spi-B and Rank) or remained unchanged (Figure 4E). These results indicate that MyD88 was involved in colitis-induced M cells differentiation.

**Figure. 4.**
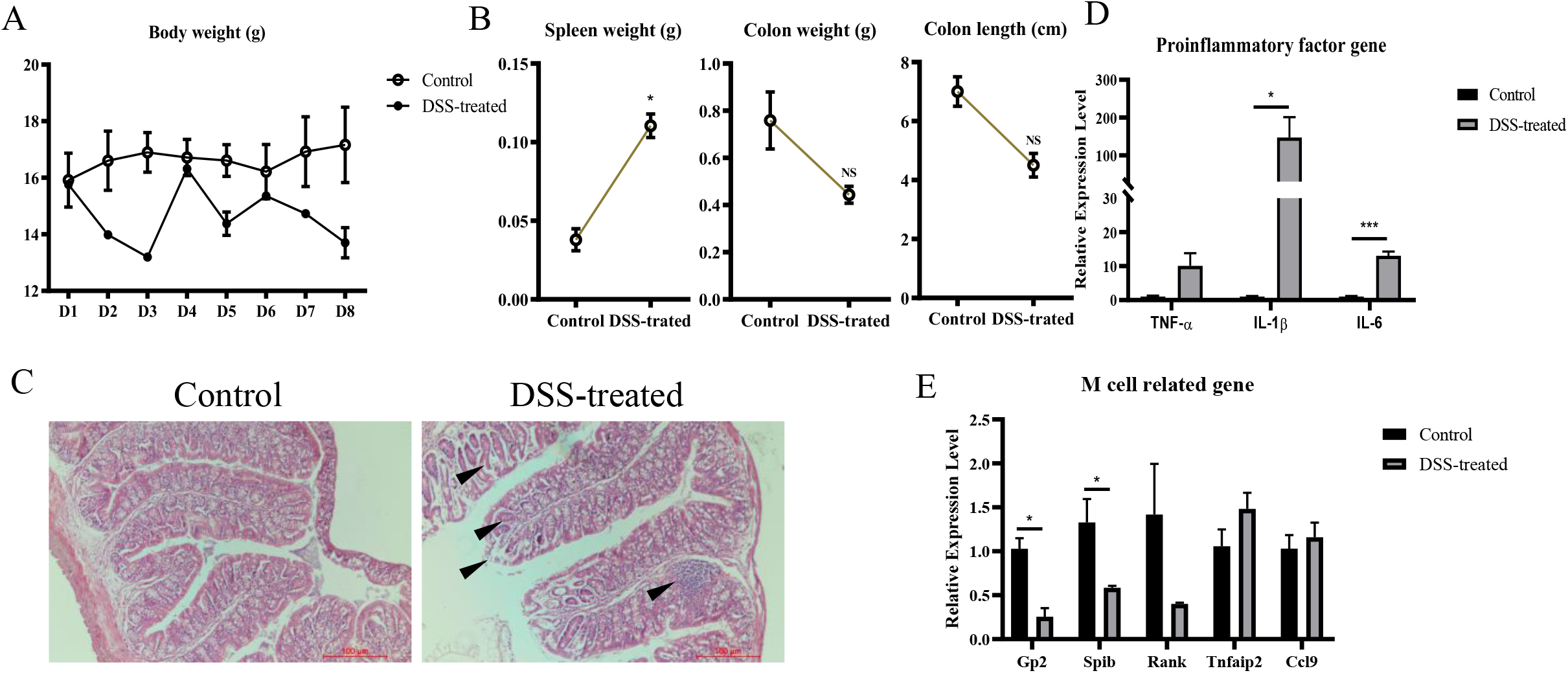
MyD88 is involved in colitis induced M cells differentiation. *MyD88*^-/-^ mice in the DSS-treated group (n=3) and control group (n=3) were allowed to intake DSS-contained or normal water for 7 days freely. The body weight of mice was monitored daily (A). The mice were sacrificed on day 8, and their spleen weight, colon weight, and colon length were measured to assess the pathological changes caused by DSS treatment (B). Total RNA from colon samples was used to detect the mRNA levels of TNF-α, IL-1β, IL-6 by RT-qPCR (C). Tissue samples from the colon were used for histological analysis to evaluate their colitis (D). Total RNA from colon samples was used to detect the mRNA levels of GP2, Spi-B, RANK, Tnfaip2, and CCL9 by RT-qPCR (E). The data were calculated using the 2^-ΔΔCt^ method. *, *P*<0.05; **, *P*<0.01; ***, *P*<0.001. The GP2 expression levels in total protein from colon samples were evaluated by Western blot (F).

### MyD88 is involved in RANKL-induced M cells differentiation

We hypothesize that MyD88 was also involved in RANKL-induced M cells differentiation. To test this hypothesis, murine intestinal organoids were prepared for further study (Figure 5A). Expression of key M cell markers was evaluated by RT-qPCR in the organoids generated from WT and *MyD88*^-/-^ mice. The results showed that M cells differentiation markers were down-regulated in *MyD88*^-/-^ mice (Figure 5B). This result supported that MyD88 was indispensable for M cells differentiation in our previous results. Then the organoids from WT mice and *MyD88*^-/-^ mice were treated with RANKL 24 hours before analysis. The results illustrated that RANKL could induce M cells differentiation in both WT mice and *MyD88*^-/-^ mice (Figure 5C), characterized as increased expression levels of GP2, Spi-B, RANK, Tnfaip2, and CCL9. However, the expression levels of M cell markers in *MyD88*^-/-^ mice were much lower than that in WT mice (Figure 5C). Taken together, these results indicate that MyD88 is also involved in M cells differentiation induced by RANKL.

**Figure. 5.**
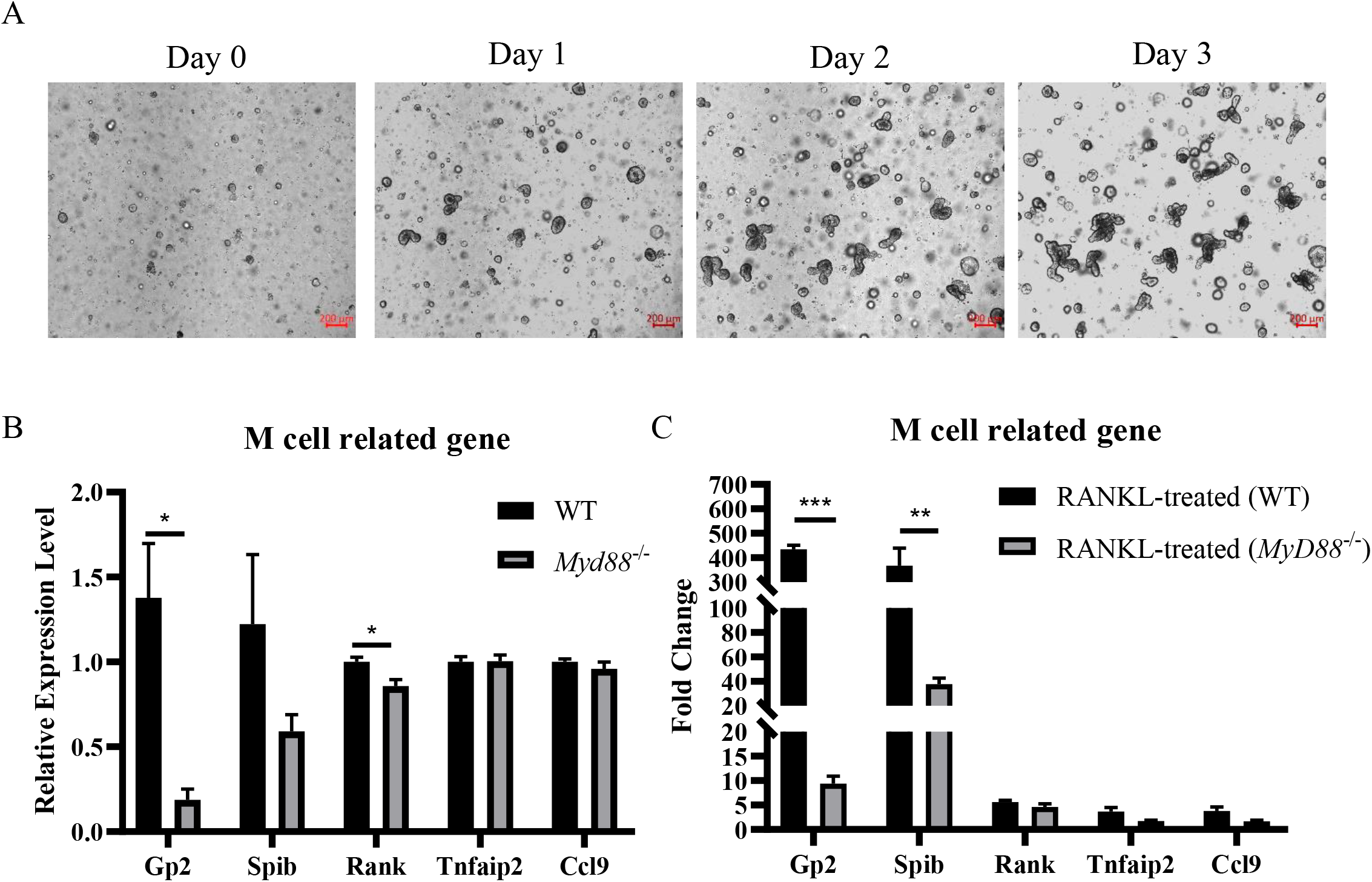
MyD88 is involved in RANKL-induced M cells differentiation. Murine intestinal organoids were established (A). Images were developed under a ZEISS Vert A1 microscope. Scale bar = 200 μm. Both WT and *MyD88*^-/-^ organoids were stimulated with human recombinant RANKL for one day (PBS was used as control) and then were collected for RNA isolation. The mRNA levels of representative M-cell related markers, GP2, Spi-B, RANK, Tnfaip2, and CCL9, in both WT and *MyD88*^-/-^ organoids, were detected by RT-qPCR (B). The fold change of these markers was compared after RANKL-treatment between WT and *MyD88*^-/-^ group (C). The data were calculated using the 2^-ΔΔCt^ method. *, *P*<0.05; **, *P*<0.01; ***, *P*<0.001.

## Discussion

M cell is a potential route for promoting drug and oral vaccine delivery. M cell-dependent antigen transport was reported to have an essential function in alleviating colitis [22]. However, the knowledge about colonic M cells differentiation is limited. The research on the differentiation of colonic M cells will help to better understand the role of M cells in maintaining intestinal homeostasis.

RANKL induces M cells differentiation. We found that GP2, used as a mature M cell marker [23], is upregulated by 2-day RANKL stimulation [24]. Our study further demonstrates that RANKL induces M cells differentiation in both WT and *MyD88*^-/-^ mice organoids, even though *MyD88*^-/-^ mice exhibit lower expression levels of M cells differentiation-related genes comparing to that in WT mice organoids.

Another study showed that gut microbiota could also regulate the differentiation of M cells [14]. As reported, *S. Typhimurium* induces the differentiation of FAE enterocytes into M cells based on a type □ effector protein SopB-dependent pathway [15]. In our study, we found that *S. choleraesuis*-induced colitis also promotes M cells differentiation. Combined with the results that DSS-induced colitis promotes M cells differentiation, we speculated that inflammation induces M cells differentiation. Moreover, we found that MyD88 plays a crucial role in both colitis- and RANKL-induced M cells differentiation. MyD88 is an essential adaptor molecule in the toll-like receptors signaling pathway, which is related to bacterial receptors [25] and intestinal microorganism homeostasis [26]. In the intestines, MyD88 also participates in regulating gut microbial ecology and maintaining homeostasis [27, 28]. Here, we found that the differentiation of M cells is attenuated in both *MyD88*^-/-^ intestinal organoids and colon of *MyD88*^-/-^ mice as compared with its WT control. These data suggested potential crosstalk between MyD88-mediated signaling and M cells differentiation.

MyD88 has been proved to play a crucial role in regulating RANKL signaling. MyD88 mediates RANKL transcription introduced by LPS and IL-1α [29]. In bone marrow stromal cells, MyD88 is also required for up-regulating RANKL transcription and down-regulating osteoprotegerin (OPG) mRNA by the TLR and IL-IR signaling [30]. As a soluble inhibitor of RANKL, OPG mediates the self-regulation of M cells differentiation in FAE [31]. These studies imply the relationship between the MyD88 mediated pathway and RANKL signaling. Hence, the deletion of MyD88 may promote the transcription of OPG and potentially inhibit M cells differentiation.

In summary, the differentiation of M cells may be regulated by a complicated signaling pathway related to immune response and gut microbiota. Here, we highlight crosstalk between inflammation response and M cells differentiation and demonstrate that MyD88 was a critical molecular for colitis- and RANKL-induced M cells differentiation. Unfortunately, due to the unsuccessful immunostaining of colonic M cells with 2 antibodies against GP2, we failed to carry out a quantitative analysis regarding the differentiation of colonic M cells in tissue. The detailed mechanism that regulates colonic M cells differentiation still needs further study.

## Abbreviations

CCL9: C-C motif ligand 9
DSS: dextran sulfate sodium
FAE: follicle-associated epithelium
GP2: glycoprotein 2
Lgr5: leucine-rich repeat-containing G-protein coupled receptor 5
M cells: microfold cells
MyD88: myeloid differentiation factor 88
NALT: nasopharynx-associated lymphoid tissue
RANK: receptor activator of NF-κB
RANKL: receptor activator of the Nuclear Factor-κB ligand
TNF-α: tumor necrosis factor-α
TRAF6: tumor necrosis factor receptor-associated factor 6

## Compliance with Ethical Statements

### Ethics approval

The care and use of animals were approved by the Institutional Animal Care and Use Committee of LVRI, Chinese Academy of Agricultural Sciences in compliance with the NIH guidelines for the care and use of laboratory animals.

### Author contributions

YL, SY, XH, NY, and GL conceived the study. YL, SY, and GL designed the experiments. YL, SY, XH, CW, JZ performed the experiments. ZJ provided *MyD88*^-/-^ mice. YL and SY analyzed data. YL, SY, LW and GL prepared the draft. GL edited the manuscript.

